# Exposure to global change and microplastics elicits an immune response in an endangered coral

**DOI:** 10.1101/2022.09.05.506609

**Authors:** Colleen B. Bove, Katharine Greene, Sharla Sugierski, Nicola G. Kriefall, Alexa K. Huzar, Annabel M. Hughes, Koty Sharp, Nicole D. Fogarty, Sarah W. Davies

## Abstract

Global change is increasing seawater temperatures and decreasing oceanic pH, driving declines of coral reefs globally. Coral ecosystems are also impacted by local stressors, including microplastics, which are ubiquitous on reefs. While the independent effects of these global and local stressors are well-documented, their interactions remain less explored. Here, we examine the independent and combined effects of global change (ocean warming and acidification) and microplastics exposures on gene expression (GE) and microbial community composition in the endangered coral *Acropora cervicornis*. Nine genotypes were fragmented and maintained in one of four experimental treatments: 1) ambient conditions (ambient seawater, no microplastics; AMB); 2) microplastics treatment (ambient seawater, microplastics; MP); 3) global change conditions (warm and acidic conditions, no microplastics; OAW); and 4) multistressor treatment (warm and acidic conditions with microplastics; OAW+MP) for 22 days, after which corals were sampled for genome-wide GE profiling and ITS and 16S metabarcoding. Overall *A. cervicornis* GE responses to all treatments were subtle; however, corals in the multistressor treatment exhibited the strongest GE responses, and genes associated with innate immunity were overrepresented in this treatment, according to gene ontology enrichment analyses. 16S analyses revealed stable microbiomes dominated by the bacterial associate *Aquarickettsia*, suggesting that these *A. cervicornis* fragments exhibited remarkably low variability in bacterial community composition. Future work should focus on functional differences across microbiomes, especially *Aquarickettsia* and viruses, in these responses. Overall, results suggest that local stressors present a unique challenge to endangered coral species under global change.

## INTRODUCTION

Anthropogenic global change represents one of the greatest scientific challenges of our time. As atmospheric concentrations of carbon dioxide (CO_2_) continue to increase, the world’s ecosystems face unprecedented challenges from the resulting warming, sea level rise, and severe extreme weather events (Rosenzweig et al., 2008). The effects of global change are being documented across ecosystems, from declines in terrestrial plant diversity (Harrison, 2020) to shifts in species distributions (Cristofari et al., 2018). However, marine environments are faced with additional threats not present in terrestrial ecosystems (*e*.*g*., sea-level rise and ocean acidification), and these threats are especially pressing for tropical coral reefs (Hoegh-Guldberg and Bruno, 2010).

Coral reef ecosystems are experiencing major declines under global change stressors, especially ocean warming that causes the breakdown of the symbiosis between the coral host and dinoflagellate algae (family Symbiodiniaceae; (Muller-Parker et al., 2015; LaJeunesse et al., 2018)). These declines are particularly evident across the Caribbean where reefs have experienced significant coral loss since the late 20th century, largely due to major disease outbreaks, overfishing leading to macroalgal blooms, and thermal stress events (Contreras-Silva et al., 2020; Bove et al., 2022b; Randazzo-Eisemann et al., 2022). In addition to warming, ocean acidification represents a significant challenge to reef-building corals that can lead to reduced growth rates, dissolution of existing reef framework, and reduced holobiont physiology and metabolic rates (Anthony et al., 2008; Aichelman et al., 2021; Cornwall et al., 2021). While responses of corals under ocean acidification and warming can vary both within and across species (Bove et al., 2021), it is clear these global stressors represent immediate threats to the future of coral reefs.

Along with changing oceanic conditions, coral reefs are susceptible to pollution via terrestrial input, which includes microplastics (Hall et al., 2015; Soares et al., 2020; Hankins et al., 2021; Oldenburg et al., 2021). Microplastics are small plastics, such as beads or fibers, smaller than 5 mm, that come from a variety of sources, including synthetic clothing and personal care products, that enter the waterways via runoff and wastewater treatment. Microplastics are ubiquitous across coastal ecosystems, and pose risk to ecosystems via adsorbed chemical pollutants and novel microbiomes (Browne et al., 2011; Rochman et al., 2019). Microplastic particles have been found within coral gastrovascular cavities, suggesting corals ingest plastics (Hall et al., 2015; Hankins et al., 2018; Rotjan et al., 2019). Active ingestion of microplastic particles can reduce coral growth (Reichert et al., 2019; Hankins et al., 2021; Huang et al., 2021) directly via blockages of coral digestive cavities (Allen et al., 2017). Further, interaction with microplastics may impact the energetic budget of the coral, altering polyp behavior and feeding, impeding gamete fertilization success, and affecting the holobiont immune system and coral disease prevalence and susceptibility (Tang et al., 2018; Berry et al., 2019; Rotjan et al., 2019; Hankins et al., 2021; Huang et al., 2021). Passive surface interactions/surface adhesion with the plastics (Reichert et al., 2018) can cause abrasion and injury, increasing coral susceptibility to infection and disease (Page and Willis, 2008; Lamb et al., 2016). While the impacts of microplastics on corals are an active area of research, little is known about how this emerging stressor interacts with other stressors, especially ocean warming and acidification.

Of the work to date examining global change stressors and microplastics, results are variable and species dependent (Axworthy and Padilla-Gamiño, 2019; Huang et al., 2021). For example, recent work found that warming led to consistent reductions in fitness-related traits in corals, while microplastics resulted in mixed responses (Reichert et al., 2021). Similarly, another study found that microplastics had no effect on Symbiodiniaceae cell density under ambient or elevated temperatures (Plafcan and Stallings, 2022), suggesting that microplastics may represent a minor stressor when compared to warming. Little is known about the combined effects of ocean warming and microplastics on corals and even less work has explored how ocean acidification may interact with microplastic pollution to impact corals. Previous work in other marine invertebrates reports impaired immunity in adult mussels (Huang et al., 2022) and altered larval development in urchins (Bertucci et al., 2022) in response to microplastics and ocean acidification, suggesting that the interaction of these stressors may exacerbate coral stress.

One way to gain a more mechanistic understanding of how an organism is responding to a stressor is to profile their gene expression in response to multiple stressors (reviewed in (Rivera et al., 2021)). Currently, gene expression responses of corals to the combined effects of global change and microplastics remain underexplored. Corals have been shown to shuffle their algal symbionts in response to a variety of stressors (Ros et al., 2021; Rodriguez-Casariego et al., 2022), representing a potential acclimation strategy for corals under stress. In addition to their obligate, intracellular photosymbionts (Symbiodiniaceae), corals maintain diverse but specific microbiomes consisting of of bacteria, archaea, fungi, viruses, and protists (Bourne et al., 2016; van Oppen and Blackall, 2019). Research has demonstrated that coral health and survival is mediated by their microbiomes (Glasl et al., 2016; Ziegler et al., 2017; Ricci et al., 2019), and that exposure to stressors is reflected in dysbiosis, or a microbiome-wide disturbance/shift in taxonomic composition (Zaneveld et al., 2016; Apprill, 2017; McDevitt-Irwin et al., 2017). Laboratory experiments have demonstrated the capacity of microplastic particles to transmit bacteria into corals via particle ingestion (Rotjan et al., 2019). Moreover, microplastics are known to harbor microbial biofilms that are taxonomically distinct (the “plastisphere”, (Zettler et al., 2013; Amaral-Zettler et al., 2020)) from those found in naturally occurring particles suspended in seawater, and bacteria associated with coral diseases have been detected on microplastic pieces (Goldstein et al., 2014; Feng et al., 2020). However, the implications of exposure to microplastics and microplastics-associated microbes on the composition and activity of the coral microbiome, including vectoring pathogens to corals, is not yet well understood.

Understanding how all of the members of the coral holobiont respond in highly endangered coral species, such as *Acropora cervicornis*, remains a top research priority given the current rates of coral decline (Miller et al., 2002; Schutte et al., 2010; Contreras-Silva et al., 2020). Caribbean *A. cervicornis* populations are experiencing dramatic declines and are the focus of major restoration efforts (Schopmeyer et al., 2017; Ware et al., 2020). Disease susceptibility in *A. cervicornis* appears to be mediated by host genotype, environment, and the microbiome (Klinges et al., 2020; Williams et al., 2022). The recently described “*Candidatus* Aquarickettsia rohweri” (hereafter, *Aquarickettsia*) (Klinges et al., 2019) is extremely widespread across Caribbean acroporids and can represent as much as 99% of the detected sequences in 16S-based studies (Klinges et al., 2020). *Aquarickettsia* appears to be parasitic to acroporids, and is likely an indicator and determinant of disease susceptibility. Conversely, taxa from the genus *Endozoicomonas* appear to be significantly and directly correlated with disease resistance and resilience (Chu and Vollmer, 2016). After bleaching/thermal stress, *Aquarickettsia* abundance declines dramatically in the host, suggesting that disease is the indirect result of *Aquarickettsia* depletion of holobiont nutritional deficiencies, providing a niche for opportunistic pathogens during/following thermal stress (Klinges et al., 2020). Understanding how the microbiomes of these important reef-building corals are involved in holobiont response to stressors is an important consideration for informed conservation, restoration, and management (Ware et al., 2020).

Here, we reared fragments of *Acropora cervicornis* under projected global change (ocean acidification and warming), microplastic pollution, and their interaction for 22 days. We assessed responses of the holobiont to these stressors via coral gene expression profiling, Symbiodiniaceae community ITS2 metabarcoding, and microbiome community 16S metabarcoding. Overall, we hypothesized that the combined stressor treatment (ocean acidification, warming, and microplastics) would elicit the strongest responses by all holobiont members given that these stressors may interact to cause cellular disruption and limit heterotrophy through microplastic ingestion.

## MATERIALS AND METHODS

### Coral collection and experimental design

In June 2020, Nova Southeastern University staff collected nine putative *Acropora cervicornis* genotypes off coastal Fort Lauderdale, FL, USA (Florida Fish and Wildlife Conservation Commission permit #SAL-19-2200A-SRP). These genotypes were maintained in outdoor tanks for one week in ambient conditions, after which they were transported to the University of North Carolina Wilmington. Fragments were acclimated to laboratory tank conditions (27.6–28.9 °C; 34–36 ppt salinity) for three months. Two weeks before the start of the experiment, four branches (∼5 cm) from each genotype were glued to ceramic plugs, allowed to recover for one week, and then acclimated to individual experimental tanks for one week.

Coral fragments were randomly assigned to one of four experimental treatments: 1) ambient treatment (AMB; 28.9 °C, 8.07 pH), 2) microplastics treatment (MP; 29.0 °C, 8.06 pH), 3) ocean acidification and warming treatment (OAW; pH: 30.5 °C, 7.94 pH), and 4) multistressor treatment (OAW+MP; 30.5 °C, 7.92 pH) (**Figure 1**; see Supplemental Table S1 for details on seawater conditions at tank and treatment levels). Ambient seawater temperature and pH were based on oceanic averages in Fort Lauderdale, FL, while OAW conditions were intermediate predictions for 2075 following IPCC data (IPCC 2019). The microplastic pollution conditions were created using two size classes of UV-fluorescent blue polyethylene microspheres (1.13 g/cc; Cospheric, LLC) with radii of 180-212 μm and 355-425 μm to ensure the corals could ingest the plastics and make it more ecologically relevant. One branch from each genotype (N=9) was placed in each treatment, resulting in a total of 36 samples. Each fragment was housed in its own non-recirculating experimental tank and thus can be considered a true biological replicate.

**Figure 1:**
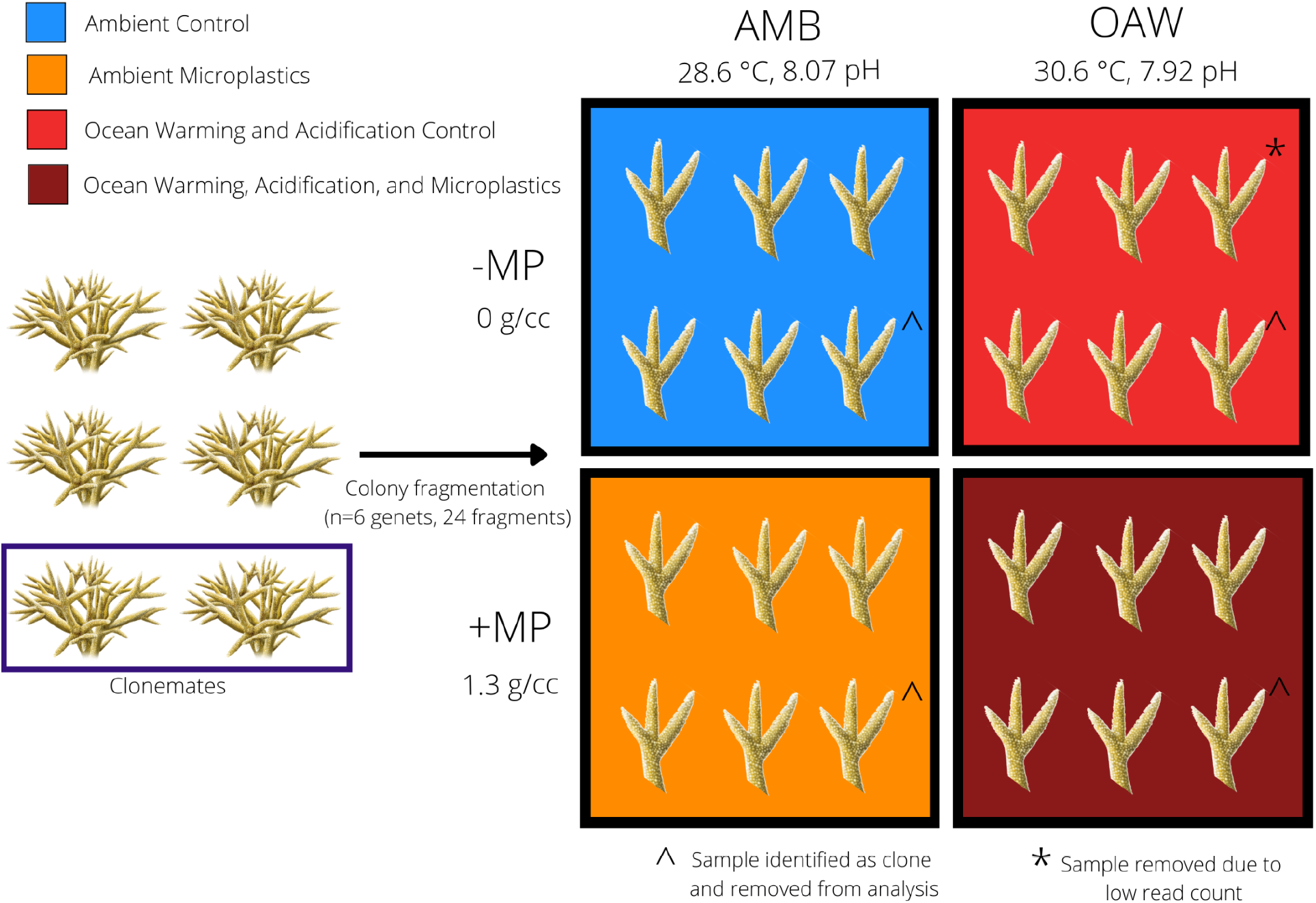
Experimental design to test the effects of ocean warming, acidification, and microplastics on the endangered coral *Acropora cervicornis*. Four fragments from each of six *A. cervicornis* genotypes that were used in downstream sequencing were assigned one of four experimental treatments: ambient conditions (AMB; blue), ocean acidification and warming (OAW; red), microplastics (MP; orange), and OAW and MP (OAW+MP; dark red). Fragments were maintained individually in isolated tanks not depicted in this diagram. The dark blue rectangle (left) showcases two corals that were identified as clones via SNP calling from TagSeq gene expression data and the ^ symbol indicates the clone pair that was removed. The asterisk indicates one gene expression library that was removed from downstream analysis due to low read counts.

Water temperature and pH were controlled using Neptune Systems’ Apex microcontrollers to modulate aquarium heaters and solenoids that bubbled CO_2_ into tanks. Fragments in OAW treatment tanks were acclimated by starting at 28.6 °C and then raising the temperature by 0.5 °C each day until 30.5 °C was achieved. Similarly, the pH in OAW treatments started at 8.055 and was reduced by 0.05 pH units each day until 7.94 was reached. Corals were maintained at these experimental conditions for 22 days while monitoring salinity, temperature, pH, and dissolved oxygen daily (Supplemental Table S1). Ammonia, nitrate, phosphate, and alkalinity were also measured daily in two randomly selected experimental tanks, one ambient and one ocean acidification and warming. Lights were on a 12:12 light:dark schedule with a PAR of 195 μmol m^2^ S^−1^ that is similar to those recorded for waters across southeast Florida (Yates et al., 2019). All fragments were fed Reef-Roids coral food following manufacturer’s instructions (2 mL aliquots) three times weekly and 30% water changes were conducted for all tanks the day after feeding. Immediately following the 3-week experiment, tissue samples (1 cm) from each fragment were placed in cryovials, flash frozen in liquid nitrogen, and maintained at −80 °C before being transported to Boston University for downstream analyses.

### RNA isolation and TagSeq library preparation

Tissue samples were crushed with a razor blade and RNA was isolated using RNAqueous kits (ThermoFisher Scientific) following manufacturer instructions with the exception of eluting in 30 μl of elution buffer. DNA was then removed using DNA-free DNA Removal kits (ThermoFisher Scientific). Of the nine genotypes, the top six (Figure 1; Table S2) with the highest concentration and quality of RNA were normalized to 15 ng/μl. Normalized samples were sent to UT Austin’s Genome Sequencing and Analysis Facility, where TagSeq libraries were prepared following Meyer et al (2011) and sequenced (single end 100 bp) on a NovaSeq 6000.

### *Identifying* Acropora cervicornis *clones*

*Acropora cervicornis* is well-known for its ability to reproduce asexually via fragmentation (Tunnicliffe, 1981). To identify potential *A. cervicornis* clones in our dataset, we called single nucleotide polymorphisms (SNPs) from TagSeq reads. Briefly, 104.64 million raw reads were generated, with individual library counts ranging from 4.17 to 6.58 million reads per sample (mean = 5.23 million reads) (Table S2). *Fastx_toolkit* removed 5’-Illumina leader sequences and poly(A)^+^ tails. Sequences <20bp in length with <90% of bases having quality cutoff scores <20 were also trimmed. In addition, because degenerate bases were incorporated during cDNA synthesis, PCR duplicates were removed from all libraries. After quality filtering 0.63 to 3.00 million reads remained (mean = 2.07 million) and these resulting quality filtered reads were mapped to the *Acropora millepora* genome (Fuller et al., 2020) using *Bowtie2*.*2*.*0* (Langmead and Salzberg, 2012) (Table S2). Resulting SAM files were converted to BAM files using *samtools* (Li et al., 2009). *ANGSD* (Korneliussen et al., 2014) calculated pairwise identity-by-state (IBS) matrices using a minIndDepth filter of five. IBS matrices were used as input to *hclust*(), which visualized relatedness of individuals and allowed for the identification and removal of clones. We note here that mapping was also conducted using the *Acropora cervicornis* transcriptome (Libro et al., 2013) and the *A. cervicornis* genome (Kitchen et al., 2019), however the transcriptome was found to contain symbiont data and the genome resulted in lower mapping efficiencies relative to the *A. millepora* genome. Given that results remained the same (*e*.*g*., clones were identified and similar overall gene expression patterns were observed), *A. millepora* has superior annotations, and Cooke *et al. (2020)* found high synteny among acroporids, we opted to move forward mapping to the *A. millepora* genome. When clones were detected, only one clonemate was maintained in downstream analyses of gene expression patterns (see Table S2).

### Read mapping, differential gene expression and Gene Ontology enrichment analysis

Reads were trimmed as described above and these reads were mapped to the *Acropora millepora* genome (Fuller et al., 2020) using *Bowtie2 (Langmead and Salzberg, 2012)*. A custom Perl script (samcount.pl) was used to count the number of reads and the resulting raw counts file was then imported into R v. 3.4.2 (R Core Team (2017)) for further analyses. Only samples representing clonemates from SNP analyses above and samples with < 400k counts were removed leaving 19 samples in downstream analyses. Differentially expressed genes (DEGs) were identified using DESeq2 v. 1.32.0 in R, with the model: *design* = ∼ *genet* + *treatment*. A contig was considered significantly differentially expressed if it had an FDR adjusted p-value < 0.1.

Data were *rlog-*normalized and effects of genotype and treatment on global gene expression profiles were tested using a PERMANOVA *(adonis()* function; *vegan* (Oksanen, 2007)) and visualized using principal components analysis (PCA) using Euclidean distances. An additional PCA was computed using only the top 1000 differentially expressed genes for experimental treatment based on raw *p*-values.

Gene expression plasticity was then calculated using the first two principal component (PC) axes as the distance between a genotype’s gene expression in its treatment condition relative to its expression in ambient conditions using a custom function (Bove, 2022). The effect of treatment on calculated distances (*i*.*e*., gene expression plasticity) was assessed using an analysis of variance (function *aov*) with treatment as a fixed effect. Genotype was not included in the model because it was accounted for in the PC distance calculations. Differences between treatment levels were tested using Tukey’s Honest Significant Differences (HSD) tests.

To determine whether global gene expression patterns showed enrichment of different gene ontology (GO) classes, the collection of scripts ‘GO_MWU’ from Wright *et al*. (2015) was used (https://github.com/z0on/GO_MWU). Here, the GO database (go.obo v.1.2) was used to test for enrichment of GO terms based on the ranked −*log* signed *p*-values of each gene. Gene ontology terms that were over-represented or under-represented were then visualized in a tree format that groups GO terms with other terms of similar function. All GO enrichment results for all treatment comparisons can be found on the github repository (https://github.com/daviessw/Acer_OAW-Microplastics); however only results of the double stressor treatment (OAW+MP) relative to ambient conditions (AMB) are described in detail here. Within this comparison, a heatmap of genes (raw *p*-value 0.10) assigned to GO terms associated with immunity was constructed using the R package *pheatmap* to illustrate gene expression patterns associated with OAW+MP.

### ITS2 and 16S metabarcoding

DNA was extracted using an RNAqueous kit (ThermoFisher Scientific) as described above, except samples were not subjected to the DNA removal step. Internal transcribed spacer region 2 (ITS2) PCR amplification was performed using *SYM_VAR_5*.*8S2* and *SYM_VAR_REV* primers (Hume et al., 2013, 2015) using the following PCR profile: 26 cycles of 95°C for 40 s, 59°C for 2 min, 72°C for 1 min and a final extension of 72°C for 7 min. A negative control was included and failed to amplify so it was not sequenced. PCR products were cleaned using the GeneJET PCR Purification kit (ThermoFisher Scientific) according to the manufacturer’s instructions with the exception of eluting in 30 μl of elution buffer. A second PCR was performed to dual-barcode samples before pooling, which was done based on the visualization of band intensity on a 1% agarose gel. After pooling, the sample was cleaned using the GeneJET PCR Purification kit (ThermoFisher Scientific) according to the manufacturer’s instructions with the exception of using 40 μl of elution buffer. 20 μl of the pool was run on a 2% SYBR Green gel, the target band was excised and placed in 30 μl of Milli-Q water overnight at 4 °C before submission for sequencing as detailed below. The V4 region of the 16S rRNA gene was amplified via PCR using Hyb515f (Parada et al., 2016) and Hyb806R (Apprill et al., 2015) primers and the following PCR profile: 30 cycles of 95 °C for 40 s, 63 °C for 2 min, 72 °C for 1 min and a final extension of 7 min. The same procedure as described above for the ITS2 samples was then followed, but with the addition of two negative controls using Milli-Q water which were included in sequencing submission. Concentrations of the ITS2 and 16S pools (via DeNovix DS-11+ Spectrophotometer) were used to combine the two pools in a 1:3 ratio, respectively. Libraries were sequenced on Illumina MiSeq (paired-end 250 bp) at Tufts Genomics Core Facility.

Demultiplexed reads were pre-processed using *bbmap* (Bushnell, 2014) to split ITS2 and 16S reads based on primers, while tossing reads that included neither primer. Resulting ITS2 reads were then analyzed by submitting paired fastq.gz files directly to *SymPortal*, which identifies specific sets of defining intragenomic ITS2 sequence variants (DIVs) to define ITS2 type profiles that are indicative of genetically differentiated Symbiodiniaceae taxa (Hume et al., 2019).

16S primers were removed using *cutadapt* (Martin, 2011), then *DADA2* (Callahan et al., 2016) was used to conduct quality filtering and inference of 3,493 amplicon sequence variants (ASVs) (see Table S3 to track reads lost through filtering). Taxonomy was assigned with *DADA2* using the *Silva* v. 138.1 database (Quast et al., 2013) and National Center for Biotechnology Information’s nucleotide database using *blast+* (Camacho et al., 2009). ASVs matching mitochondria, chloroplasts, or non-bacterial kingdoms were removed (216 total) and 5 ASVs were removed based on negative controls as contaminants (*Decontam*; (Davis et al., 2018)). Cleaned counts were rarefied to 9,409 using *vegan* (Oksanen, 2007) and trimmed using *MCMC*.*OTU* (Wright et al., 2015) to remove ASVs present in less than 0.01% of counts, resulting in 260 ASVs across all 36 samples.

Beta diversity of the 16S trimmed counts based on treatment and genotype was assessed using a PCoA on Bray–Curtis dissimilarity (*Phyloseq*; (McMurdie and Holmes, 2013)) and a PERMANOVA (*vegan*; (Oksanen, 2007)). We then calculated alpha diversity (Shannon index, Simpson’s index, ASV richness, and evenness) of the rarefied counts using Phyloseq (function *estimate_richness*(); (McMurdie and Holmes, 2013)). Finally, to test for differences in background 16S communities, we removed all ASVs from the dominant genus MD3-55 (*Candidatus* Aquarickettsia rohweri, referred now as “*Aquarickettsia*”; (McMurdie and Holmes, 2013; Klinges et al., 2019)) and performed the same beta and alpha diversity assessments. All ITS2 and 16S data, figures, and analyses were completed in R (version 3.6.3; R Core Team (2020)) and can be found on GitHub (https://github.com/seabove7/acer_microplastics).

## RESULTS

### Presence of clones

A total of 4,018 SNPs were identified and of the six putative genotypes that were sequenced, one pair of clones was identified (G10a and G12) (Supplemental Figure 1). Thus, genotype G12 samples from each of the four experimental treatments were removed to retain only one representative clone (genotype G10a) in each treatment (Table S2).

### Symbiodiniaceae community compositions

Raw ITS2 counts before submission to *SymPortal* ranged from 4,684 to 613,872 per sample, with a mean of 169,246. After classification by *SymPortal*, ITS2 counts ranged from 730 to 121,515 per sample across all 36 samples with mean 32,774 counts. All coral fragments hosted *Symbiodinium ‘fitti’* (ITS2 type A3) with 44% (n=16) hosting small background amounts (less than 5%) of *Breviolum minutum* (ITS2 type B2).

### Overall differential expression responses to experimental treatments

A total of 20 samples were sequenced, which resulted in a total of 104.6 million reads, 41.4 million of which remained after trimming, and 13.9 million of which mapped successfully to the *A. millepora* reference genome. One sample - genotype G2, from the OAW treatment - was removed from the dataset due to low counts (166,828 total; Supplemental Table 1). Mean counts across samples after outlier removal was 661,358.

Pairwise comparisons between each stressor (MP, OAW, OAW+MP) and ambient conditions (AMB) showcased that individuals in the the double stressor (OAW + MP) treatment had the highest number of differentially expressed genes (DEGs) (79 up, 58 down), followed by microplastics (48 up, 9 down) and then OAW (18 up, 18 down) with relatively few genes overlapped between treatments (pfdr<0.10, Supplemental Figure 2). A principal component analysis (PCA) on all *rlog-*transformed counts found that there was no overall effect of treatment on *A. cervicornis* gene expression; however, there was a significant effect of genotype on overall expression patterns (Figure 3A; pGENET<0.001). A PCA of the top 1,000 DEGs showcased stronger clustering by experimental treatment and the effect of genotype remained significant (Figure 3B; pGENET<0.001, pTREATMENT<0.001). This second PCA also showcased that the double stressor (OAW+MP) had the strongest effect on gene expression, which corroborated the numbers of observed DEGs. A gene expression plasticity analysis confirmed this pattern, which showcased a significant effect of treatment on gene expression plasticity (pTREATMENT=0.041), with the OAW+MP treatment exhibiting the highest plasticity (Figure 3C). However, these differences in plasticity between corals in the OAW+MP treatment relative to corals in the OAW and MP treatments were only marginally significant after multiple test correction (OAW+MP∼MP, p=0.07; OAW+MP∼OAW, p=0.06).

**Figure 2:**
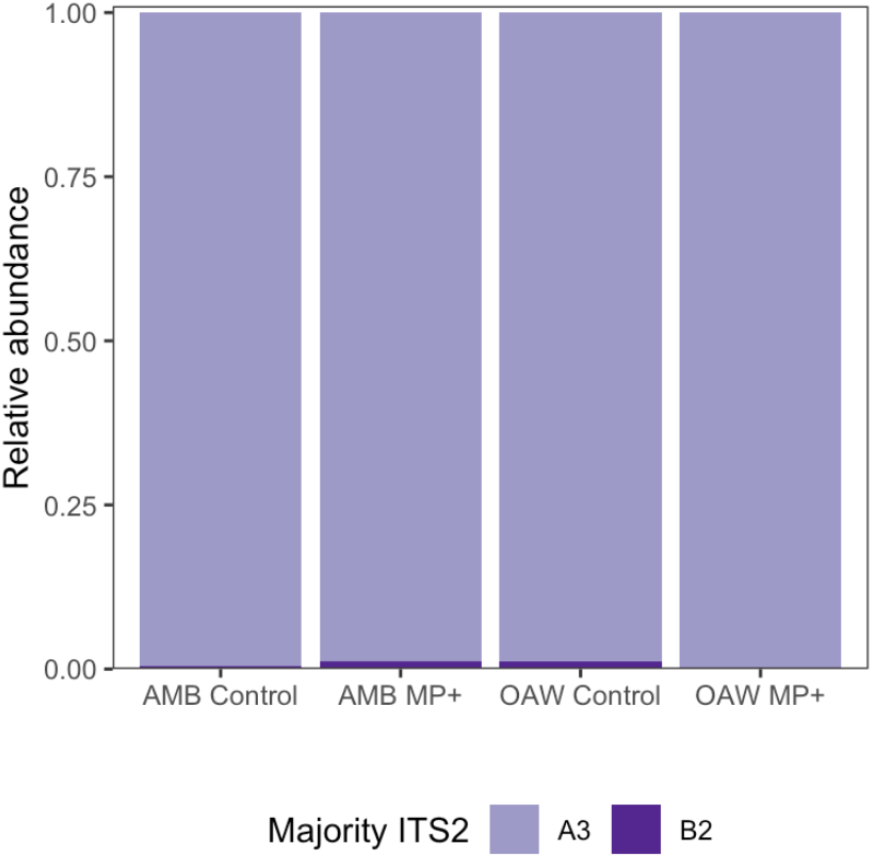
Relative abundance of major ITS2 types grouped by treatment (n = 9 corals per treatment). Light purple represents *Symbiodinium ‘fitti’* (A3) and dark purple represents *Breviolum minutum* (B2).

**Figure 3:**
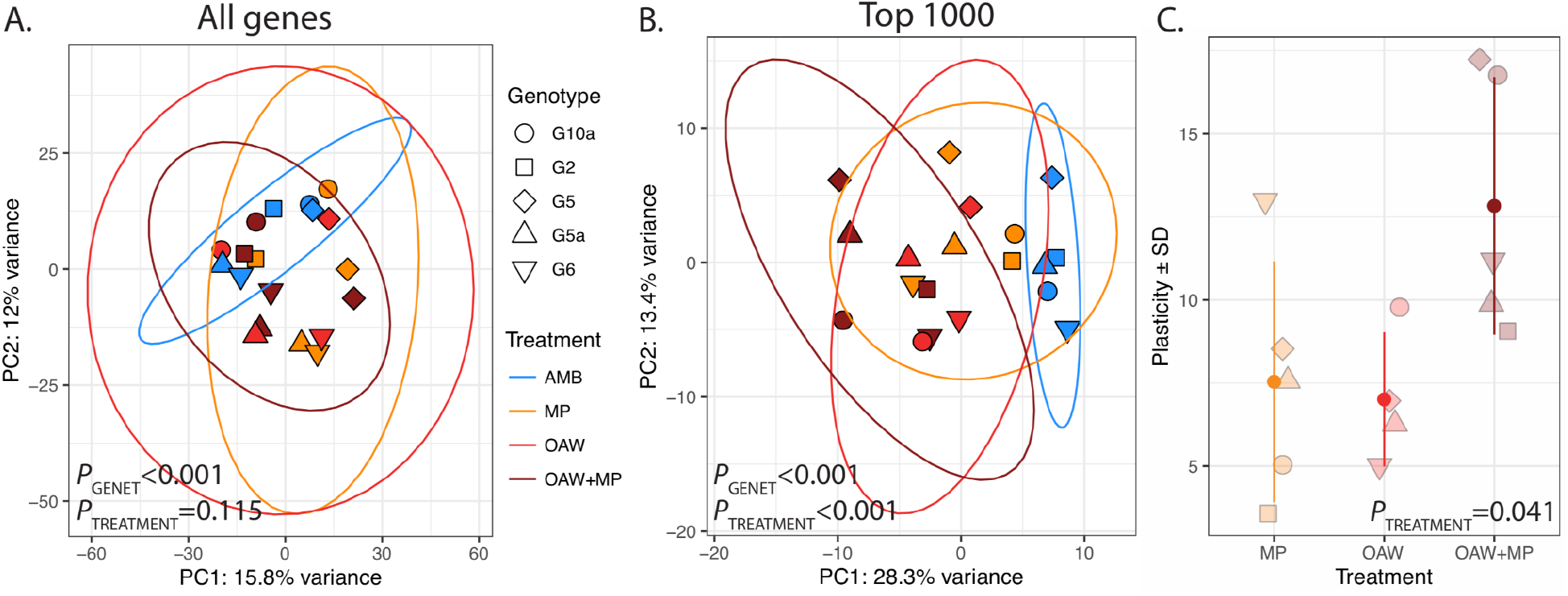
Transcriptomic responses to local and global stressors in *Acropora cervicornis*. **(A)** Principal component analysis (PCA) of *rlog*-transformed counts for all genes and, (**B**) the top 1000 differentially expressed genes by treatment in *DESeq2*. PERMANOVA results for the effect of genetic background (genet) and treatment are included. **(C)** Gene expression (GE) plasticity +/-standard deviation (SD) estimates calculated from PC distances from (**B**) where each point is the distance that genotype moved from its ambient fragment (AMB) in PC space.

### Functional enrichment in response to single stressors

Gene ontology enrichment analysis between microplastics (MP) relative to ambient conditions (AMB) revealed underrepresentation of *oxidoreductase* (GO: 0016491), suggesting down regulation of stress response in corals in MP treatment. In addition, overrepresentation of GO terms associated with ribosomal processes was observed including *structural constituent of ribosome* (GO:0003735) and *large ribosomal subunit* (GO:0015934), suggesting upregulation of terms associated with increased growth under the presence of microplastics.

In response to OAW, we observed an enrichment of *regulation of defense response to virus* (GO:0050688), terms associated with amino acid catabolism (*aromatic amino acid family catabolic process* (GO:0009074), *cellular amino acid catabolic process* (GO:0009063) and *alpha-amino acid catabolic process* (GO:1901606)), and two terms associated with stress response (*chaperone-mediated protein folding* (GO:0061077) and *unfolded protein binding* (GO:0051082)). In addition, we observed downregulation of processes associated with sensory detection including *detection of stimulus* (GO:0051606), *sensory perception* (GO:0007600), and *cognition* (GO:0050890). Lastly, several GO terms associated with synapse were down regulated including *synapse* (GO:0045202), *synaptic membrane* (GO:0097060), and *cell junction* (GO:0030054).

### Enrichment of immune-related functions under multiple stressors

Gene ontology enrichment analysis between our double stressor (OAW+MP) relative to ambient conditions (AMB) revealed many significantly enriched GO terms within ‘Biological processes’, of which the OAW+MP-enriched categories were dominated by GO terms associated with immunity (green box, Supplemental Figure 3). These enriched GO terms included: *regulation of defense response to virus by host* (GO:0050691), activation of innate immune response (GO:0002218), *regulation of immune effector process* (GO:0002697), *NIK/NF-kappaB signaling* (GO:0038061), *regulation of cytokine production* (GO:0001819), suggesting that the combined stressor elicited an upregulation of innate immunity in the coral host. When a heatmap was made of all genes belonging to these 30 GO terms that met our alpha cut-off (unadjusted *p*-value < 0.10), strong differences in gene expression were observed between corals in OAW+MP relative to those in AMB treatments (Figure 4). Of particular interest, many classic stress response genes were upregulated in OAW+MP relative to AMB including: superoxide dismutase, two heat shock proteins, several ubiquitin genes, tumor necrosis factors/receptors, a proto-oncogene, and several genes associated with apoptosis (bcl2-like 1) and response to cytokines (p38 map kinase).

**Figure 4:**
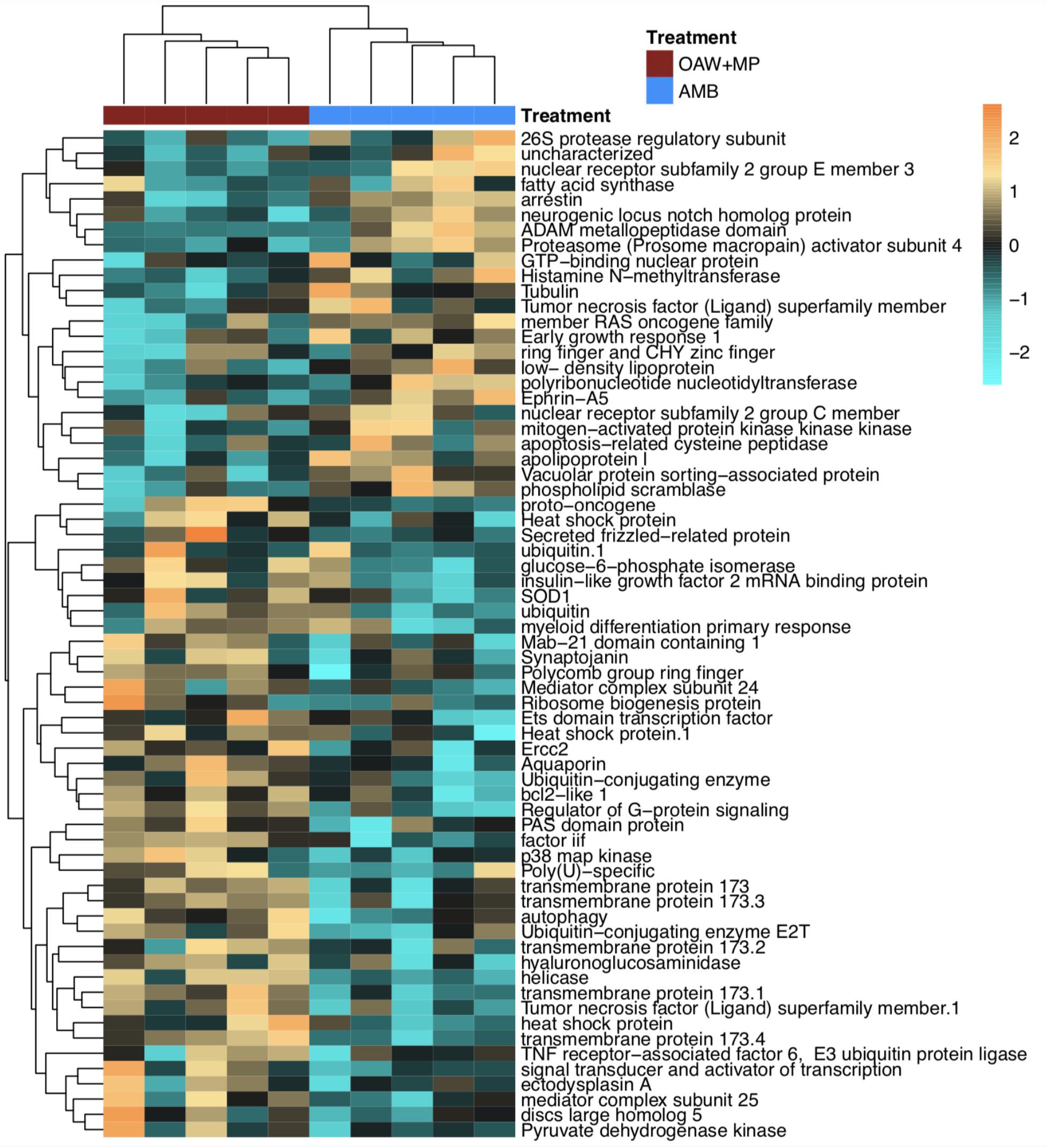
Differentially expressed genes (DEGs, unadjusted *p*-value < 0.10) with annotations associated with immunity gene ontology (GO) terms from Supplemental Figure 3 (green box). Heatmap showing annotated immunity genes where each row is a gene and each column is a unique gene expression sample. The color scale is in log2 (fold change relative to the gene’s mean) and genes and samples are clustered hierarchically based on Pearson’s correlation of their expression across samples. Colored blocks indicate treatment. Hierarchical clustering of libraries (columns) demonstrates strong differences in gene expression of immunity genes by treatment condition.

### Acropora cervicornis *microbiomes dominated by order Rickettsiales*

Cleaned 16S reads before trimming and rarefaction averaged 42,577 ± 25,010 (±SD) per sample across all *A. cervicornis* samples with a minimum of 9,409 (Sample 11 A - 1) and a maximum of 107,941 (Sample 10a C - 1). Fragments were dominated by taxa from the phylum Proteobacteria (Figure 5A), which was dominated by a single ASV (Genus MD3-55; “*Candidatus* Aquarickettsia rohweri”) within the order Rickettsiales (Figure 5B; see Supplemental Figure 4). This dominant ASV was a 100% match to the published 16S rRNA sequence for the putative bacterial parasite “*Ca*. Aquarickettsia rohweri” (hereafter, *Aquarickettsia*) (Klinges et al., 2019) that is common in *A. cervicornis* across the Caribbean (Godoy-Vitorino et al., 2017; Williams et al., 2022).

**Figure 5:**
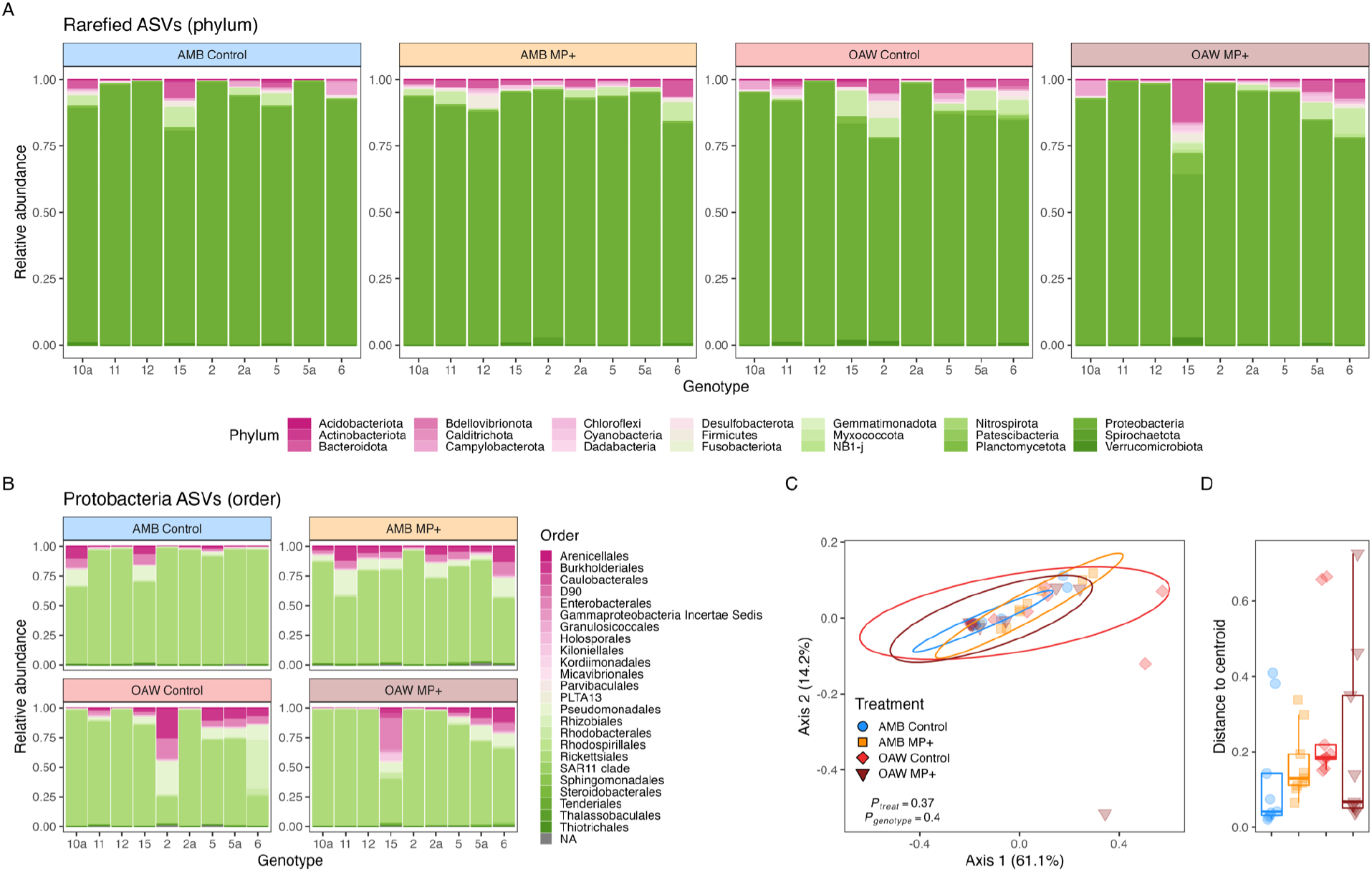
*Acropora cervicornis* bacterial (16S) relative abundance across fragments and experimental treatments. Diversity at the phylum level (**A**) depicts dominance of all samples by taxa from Proteobacteria and this (**B**) phylum was dominated by the order Rickettsiales in most samples. Bacterial diversity color was assigned alphabetically, not based on abundance. Beta diversity was visualized through (**C**) multivariate ordination plots (PCoA) of between-sample Bray–Curtis dissimilarity of rarefied ASVs of all taxa and (**D**) distance to treatment centroids. PCoA ellipses depict 95% confidence intervals and p-values indicate significance of treatment and genotype. Treatment is depicted by color in all panels: blue = ambient conditions (AMB), orange = ocean acidification and warming (OAW), red = microplastics (MP), and dark red = OAW and MP (OAW+MP).

Beta diversity analyses did not identify any statistical differences across samples based experimental treatment or genotype (Figure 5C and Table S4). Similarly, alpha diversity was indistinguishable across treatments (Supplemental Figure 5 and Table S5). After removing *Aquarickettsia* from all samples to assess background bacterial communities, there were still no differences in alpha or beta diversity across treatments or genotypes (Supplemental Figure 6 and Table S6).

## DISCUSSION

### *Microplastics alone do not drive strong gene expression responses in* Acropora cervicornis

Tropical reef-building coral ecosystems are facing extraordinary challenges from both local and global stressors that are altering their function (Eddy et al., 2021). On the local scale, pollution in the form of microplastics may threaten the health and physiology of reef-building corals (Soares et al., 2020; Nanthini devi et al., 2022). While there is evidence that corals ingest microplastics, which can lead to a variety of physiological responses (Hall et al., 2015; Rotjan et al., 2019), here we only detected a muted gene expression response in *A. cervicornis* exposed to microplastics. In fact, overall gene expression profiles were indistinguishable between the ambient and microplastics treatments, and an effect of microplastics was only apparent when looking at the expression of top 1,000 genes. This result corroborates previous work reporting negligible effects of microplastics on coral physiology, bleaching susceptibility, and mortality (Reichert et al., 2021; Plafcan and Stallings, 2022) and supports the hypothesis that corals that rely more heavily on heterotrophy may be more impacted by this stressor.

Recent studies have considered the physiological impact of microplastics on tropical corals at the molecular level. These studies, conducted on the tropical coral *Pocillopora damicornis* (Tang et al., 2018) and the habitat-forming octocoral *Corallium rubrum* (Corinaldesi et al., 2021), report evidence of elevated stress response after exposure to microplastics in as little as a few hours. In contrast, we identified an enrichment of GO terms associated with growth (structural constituent of ribosome, large ribosomal subunit) and an underrepresentation of terms associated with stress response (oxidoreductase). These contrasting results may be due to experimental design differences (microplastic concentration and size, duration, etc.); however, it is more likely due to species-specific responses (Hankins et al., 2021; Mendrik et al., 2021). Previous work on *A. cervicornis* suggests that exposure to microplastics in the presence of warming does not impact bleaching, likely because this species may not ingest the microplastics under otherwise ideal conditions (Plafcan and Stallings, 2022). Instead, *A. cervicornis* relies on photosynthetically-derived carbon from its algal symbionts as the primary source of nutrition instead of heterotrophy, thus avoiding the potential pitfalls of microplastics ingestion observed in other aggressively feeding coral species (Reichert et al., 2019; Rotjan et al., 2019; Hankins et al., 2021; Mendrik et al., 2021).

While these findings suggest that microplastics exposure alone does not elicit a severe stress response in *A. cervicornis*, it is unclear whether prolonged exposures may eventually impact the host (Hankins et al., 2021) and their algal symbionts (Lanctôt et al., 2020; Ripken et al., 2020). Further, microplastic pollution is occurring alongside global stressors (Soares et al., 2020), including widespread shifts in the microbial landscape of seawater (Cavicchioli et al., 2019) and increased prevalence of pathogens and disease across the Caribbean (van Woesik and Randall, 2017). Because microplastics have been shown to be a vehicle for microbial transmission into a corals (Rotjan et al., 2019), there is emerging concern for microplastics pollution in combination with increased pathogen exposure that has long been a predicted component of global change (Cavicchioli et al., 2019).

### Global change treatment elicits subtle stress signatures

Along with local concerns, global stressors, such as ocean acidification and warming, are contributing to the wide-scale degradation of coral reef ecosystems. Like its response to microplastics, *A. cervicornis* in the ocean acidification and warming (OAW) treatment exhibited similar overall gene expression profiles to those in the ambient treatment and only gene expression profiles of the top 1,000 genes resulted in a clear response. This subtle response to OAW in *A. cervicornis* is surprising since this species is known to be sensitive to these global stressors (Enochs et al., 2014; Towle et al., 2015; Kaufman et al., 2021; Muller et al., 2021). These contrasting results may be due to different experimental design considerations across studies (Bove et al., 2020), especially given that the heat stress temperature employed here (∼30.5 °C) was lower than most other warming studies. However, there is also significant genetic variation within *A. cervicornis* (Drury et al., 2017; Million et al., 2022), thus it is also possible that the genotypes used in this study were particularly resistant. Alternatively, these corals are already exposed to warmer seawater conditions where they were collected compared to other Caribbean reefs (Muñiz-Castillo et al., 2019; Bove et al., 2022b), suggesting the potential for acclimatization to elevated temperatures.

While we only observed minor gene expression differences in *A. cervicornis* reared in the OAW treatment, these shifts were associated with an enrichment of stress-related GO terms. For example, corals in OAW treatment exhibited enrichment of GO terms linked to protein folding and protein binding, which have been reported previously in heat-stressed corals *in situ* (Ip et al., 2022). This pattern was accompanied by enrichment of terms associated with amino acid catabolism, suggesting *A. cervicornis* may have mediated physiological stress through protein catabolism (Davies et al., 2016; Aguilar et al., 2019; Rädecker et al., 2021). While we did not assess host or symbiont physiological traits along with these gene expression profiles, it is likely that the growth and/or energy reserves may have been impacted as a result of treatment conditions (Aichelman et al., 2021; Scucchia et al., 2021; Bove et al., 2022a).

Most interestingly, we identified enrichment of GO terms associated with defense against viruses. Previous work has suggested that corals become more susceptible to infection when water temperatures increase (Bruno et al., 2007). Indeed, patterns of Caribbean coral disease have become more prevalent in recent decades (Randazzo-Eisemann et al., 2022) and population declines of *A. cervicornis* have been specifically linked to increases in disease associated with seawater temperature increases (Muller et al., 2018; Goergen et al., 2019). While the temperature stress applied here was not high enough to induce a classic heat stress response, it suggests that even subtle increases in temperature may make this species of coral more susceptible to infection.

### Global change stressors interact with microplastics to invoke an immune response

Ecological stressors can interact to pose further threats to marine organisms that may not be accounted for when assessing responses to stressors independently (Darling and Côté, 2008; Ellis et al., 2019). Indeed, our multistressor treatment (ocean acidification, warming, and microplastics; OAW+MP) resulted in the strongest gene expression response, suggesting that microplastic pollution may interact with ocean acidification and warming to elicit a more severe molecular response. This strong response to multiple stressors is commonly observed in coral (Coles and Jokiel, 1978; Reynaud et al., 2003; Courtial et al., 2017; Muller et al., 2021) and has been reported previously in studies assessing gene expression as well (Ogawa et al., 2013).

Here, we identified GO terms enriched in the multi stressor treatment related to response to viruses and innate immunity, indicating these stressors may invoke an immune response by the coral. Interestingly, a similar response was observed in OAW alone, however, to a lesser extent, suggesting that the presence of microplastics magnified this stress response. While no studies to date have assessed gene expression responses in corals to the stressors together tested here, previous work in the marine mussel, *Mytilus coruscus*, exposed to ocean acidification and microplastics identified signals of a repressed immunity that may lead to higher disease susceptibility (Huang et al., 2022). Further, gene expression patterns consistent with an upregulation of immune stress response were reported in the brine shrimp *Artemia franciscana* exposed to microplastics and warming (Han et al., 2021). This is particularly interesting because *Artemia* is also frequently used as a nutrition source in coral experiments, including microplastic ingestion experiments (Hall et al., 2015; Axworthy and Padilla-Gamiño, 2019; Lanctôt et al., 2020), potentially altering the nutritional value of the *Artemia* and impacting coral health. Overall, these results suggest that the combination of global change stressors and microplastics will likely lead to suppressed immunity in corals in tandem with higher disease transmission associated with warming (Randazzo-Eisemann et al., 2022) and microplastic pollution (Rochman et al., 2019; Rotjan et al., 2019).

### Acropora cervicornis *host stable microbiome communities across treatments*

Surprisingly similar microbiome compositions were observed across the four treatment conditions. All of the tested individuals were high-*Aquarickettsia*, ranging from 17.5% up to 98.8% of the total sequence reads. This sequence dominance in *A. cervicornis* microbiomes is not uncommon and has been documented in this species in the Florida Keys and throughout the Caribbean (Baker et al., 2022), although previous work has also showcased variation among individual genotypes (Aguirre et al., 2022). Even when we removed *Aquarickettsia*-affiliated ASVs from analysis, microbiome composition did not significantly differ across treatments. Importantly, the microbiome composition of these corals before collection from the wild is unknown, therefore we cannot determine if the universally high *Aquarickettsia* levels were a result of captivity muting microbiome signatures (Galand et al., 2018). Future work leveraging experiments on individuals previously identified as high-*Aquarickettsia* and low-*Aquarickettsia* (*e*.*g*., (Aguirre et al., 2022)) may clarify differential responses to stressors, especially since low-*Aquarickettsia* individuals may be more resilient to disease relative to high-*Aquarickettsia* individuals (Klinges et al., 2022).

Genes implicated in the coral immune response were upregulated in response to the OAW and multistressor treatment, suggesting that some sort of external stimulus triggered this response. Microbiome taxonomic profiling did not reveal significant differences in microbiomes across treatments, so we suggest that further research should investigate the transcriptomic and metabolomic responses of the bacterial and viral fractions across these stressors. For example, it is possible that while *Aquarickettsia* remained dominant, it may have differentially regulated genes in response to OAW conditions and/or OAW+MP. Alternatively, the immune response may reflect host response to physical injury/insult by contact with the plastics (van de Water et al., 2015), but not necessarily microbial infection.

The evidence of a host immune response without a corresponding bacterial community composition shift in the OAW and multistressor (OAW+MP) treatments suggests that the observed response was not due to changing bacterial composition. Another potential explanation for this is a host response to viruses. Indeed, GO categories consistent with a viral response were overrepresented in OAW+MP treatments (*i*.*e*., cellular response to virus). Coral viromes have been implicated in directing holobiont response to environmental stress (Thurber and Correa, 2011; Leruste et al., 2012; Nguyen-Kim et al., 2015; Thurber et al., 2017). It is thought that viruses may play a role in coral bleaching under thermal stress through host lysis of algal symbiont cells that precedes bleaching signals (Lawrence et al., 2014; Grupstra et al., 2022). Additionally, previous work has demonstrated increased expression of anti-viral transcripts in hosted algal symbionts (*Cladocopium*, formerly Clade C) in response to thermal stress (Levin et al., 2017). Taken together, it is possible that the *A. cervicornis* immune response may suggest an early warning of bleaching that precedes the classic environmental stress response (Barshis et al., 2013; Dixon et al., 2020).

There is also evidence that microplastics and the microbial plastisphere can facilitate survival of certain viruses (Moresco et al., 2021). It is plausible that increased levels of virus-coral interactions could elicit a host immune response, if not also facilitate viral infection of coral cells (Bowley et al., 2021; Loiseau and Sorci, 2022). We recommend that future work investigating the effects of microplastic pollution on reef-building corals, especially in combination with global change stressors, specifically target the viral dynamics and diversity in the seawater and associated with the holobiont. Additionally, assessing the microbiome and virome at several time points of stress challenge, along with host gene expression and metabolomes, will be crucial for understanding the timeline and underlying causes of potential assemblage shifts that inform holobiont phenotypes. Overall, our results highlight the importance of applying multiomic approaches to better understand the consequences of local and global stressors on reef-building corals.

## Acknowledgements

All computational work was conducted on Boston University’s Shared Computing Cluster (SCC). We thank the numerous UNCW undergraduate students and graduate students, Chad Campbell, Reanna Jeanes, and Bryce Corbett for assisting with the experimental portion. Thank you to Nova Southeastern University staff, particularly Dave Gilliam and his lab for collecting the coral and Joana Figueiredo and her lab for quarantining the corals used in this experiment. Thank you to Florida Fish and Wildlife Conservation Commission for providing permit #SAL-19-2200A-SRP for this collection.

## Data Availability

Raw reads for all 24 gene expression samples (6 genotypes x 4 experimental treatments) are available on the NCBI Short Read Archive (SRA) under BioProject number PRJNAXXXX. ITS2 and 16S metabarcoding data for all 36 experimental samples are also available on NCBI SRA under BioProject numbers PRJNAXX and PRJNAXX, respectively. All other scripts and code required to generate gene expression results described here are hosted at https://github.com/daviessw/Acer_OAW-Microplastics (Zenodo DOI: XXX) and the ITS2 and 16S materials are hosted here https://github.com/seabove7/acer_microplastics (Zenodo DOI: XXX). Metadata for samples and all other information and figures referenced in the text is supplied in the supplementary materials or main manuscript.

## Author contributions

SS, NF, and SWD conceived the research, SS and NF conducted the experiment, KG, NGK, AKH, AMH, and CBB performed molecular lab work, KG, NGK, and SWD completed the gene expression analyses, CBB, AKH, and AMH completed the 16S and ITS2 analyses, CBB and KS interpreted the 16S analyses, and CBB, KG, and SWD wrote the manuscript with help from all coauthors. Funding acquisition, project administration, and resources was led by SWD, NF, and SS.

