## Supplemental Materials for "Exposure to global change and microplastics elicits an immune response in an endangered coral"

Colleen B. Bove[1*], Katharine Greene[1], Sharla Sugierski[2], Nicola G. Kriefall[1], Alexa K. Huzar[1], Annabel M. Hughes[1], Koty Sharp[3], Nicole D. Fogarty[2], Sarah W. Davies[1*]

[1]Department of Biology, Boston University, Boston, MA USA
[2]Department of Biology and Marine Biology, Center for Marine Science, University of North Carolina Wilmington, Wilmington, NC USA
[3]Department of Biology, Marine Biology, and Environmental Science, Roger Williams University, Bristol, RI, USA

*co-corresponding authors:
Sarah W. Davies;
Colleen B. Bove;

#### Supplemental Figures

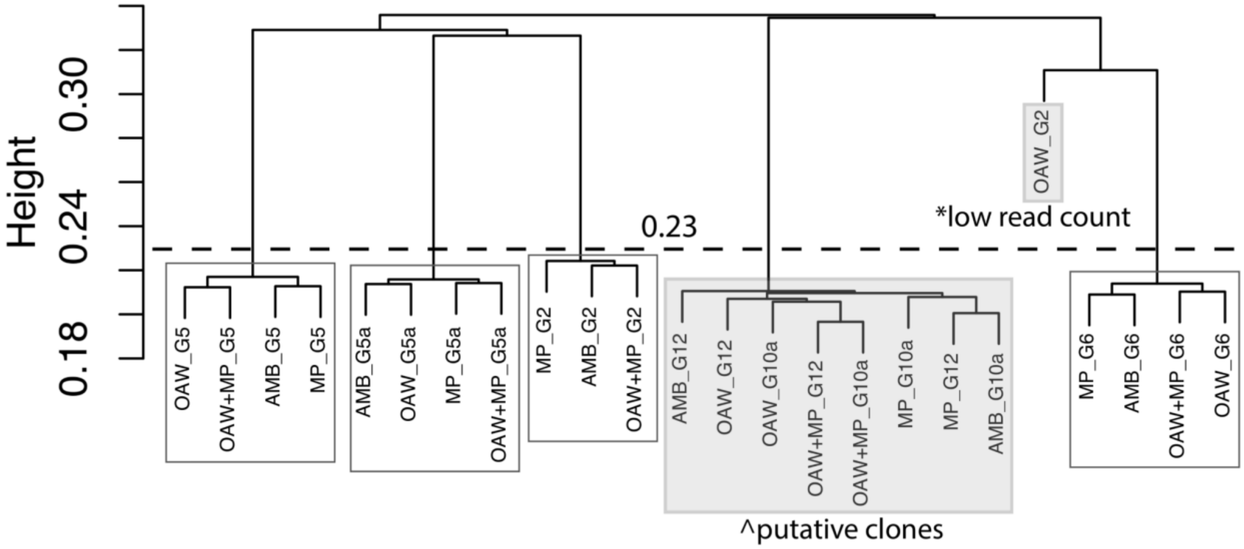


**Supplemental Figure 1**: Identity by state (IBS) dendrogram highlighting the presence of putative clones in experimental *Acropora cervicornis*. Cluster dendrogram of all samples. The gray box highlights one clonal pair (^ G12 and G10a), which occur at the same height as other technical replicates. The dashed line at height = 0.23 represents the cutoff for clonal assignment. The asterisk represents a sample that had low read count and was removed from downstream analyses.

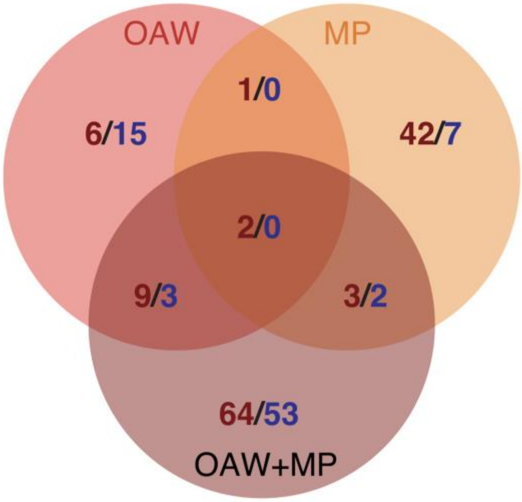


**Supplemental Figure 2**: Venn diagram of differentially expressed genes ($P_{FDR}$=0.10) with respect to experimental treatment relative to control conditions (OAW=ocean acidification and warming, MP=microplastics, OAW+MP=ocean warming and microplastics). Red numbers indicate up-regulated genes and blue numbers indicate down-regulated genes.

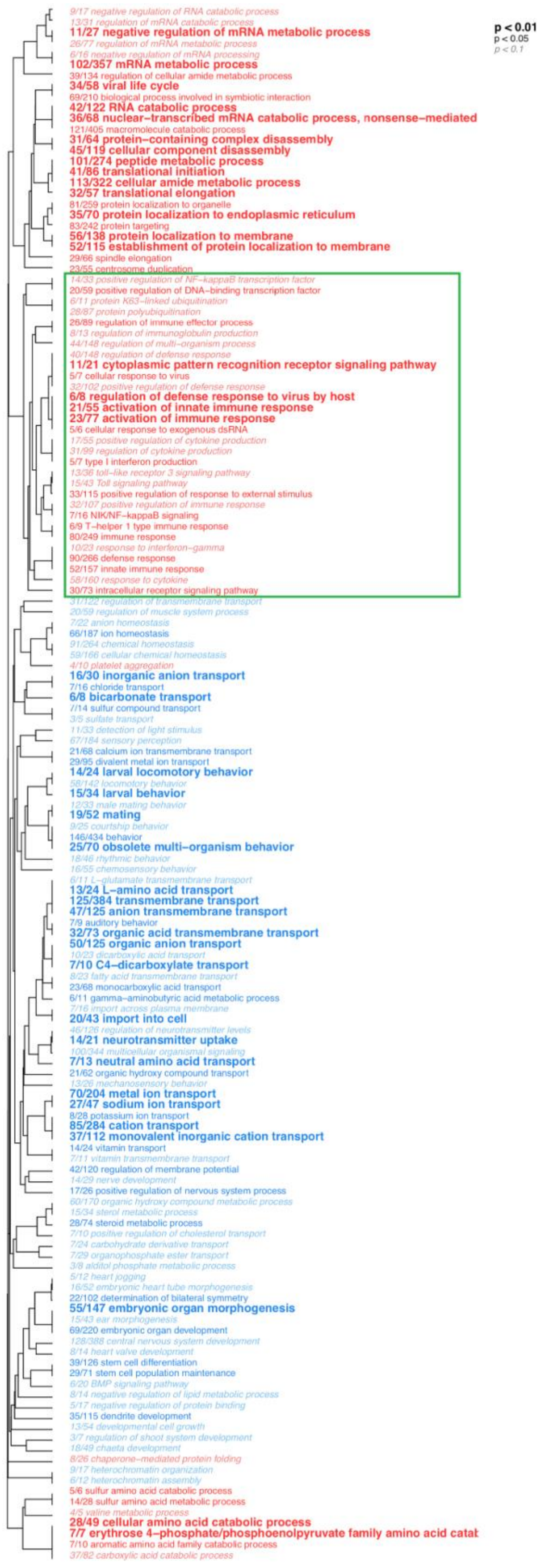

p < 0.01
p < 0.05
p < 0.1
9/17 negative regulation of RNA catabolic process
13/31 regulation of mRNA catabolic process
11/27 negative regulation of mRNA metabolic process
26/77 regulation of mRNA metabolic process
6/16 negative regulation of mRNA processing
102/357 mRNA metabolic process
39/134 regulation of cellular amide metabolic process
34/58 viral life cycle
69/210 biological process involved in symbiotic interaction
42/122 RNA catabolic process
36/68 nuclear–transcribed mRNA catabolic process, nonsense–mediated
121/405 macromolecule catabolic process
31/64 protein–containing complex disassembly
45/119 cellular component disassembly
101/274 peptide metabolic process
41/86 translational initiation
113/322 cellular amide metabolic process
32/57 translational elongation
81/259 protein localization to organelle
35/70 protein localization to endoplasmic reticulum
83/242 protein targeting
56/138 protein localization to membrane
52/115 establishment of protein localization to membrane
29/66 spindle elongation
23/55 centrosome duplication
14/33 positive regulation of NF–kappaB transcription factor
20/59 positive regulation of DNA–binding transcription factor
6/11 protein K63–linked ubiquitination
28/87 protein polyubiquitination
26/89 regulation of immune effector process
8/13 regulation of immunoglobulin production
44/148 regulation of multi–organism process
40/148 regulation of defense response
11/21 cytoplasmic pattern recognition receptor signaling pathway
5/7 cellular response to virus
32/102 positive regulation of defense response
6/8 regulation of defense response to virus by host
21/55 activation of innate immune response
23/77 activation of immune response
5/6 cellular response to exogenous dsRNA
17/55 positive regulation of cytokine production
31/99 regulation of cytokine production
5/7 type I interferon production
13/36 toll–like receptor 3 signaling pathway
15/43 Toll signaling pathway
33/115 positive regulation of response to external stimulus
32/107 positive regulation of immune response
7/16 NIK/NF–kappaB signaling
6/9 T–helper 1 type immune response
80/249 immune response
10/23 response to interferon–gamma
90/266 defense response
52/157 innate immune response
58/160 response to cytokine
30/73 intracellular receptor signaling pathway
31/122 regulation of transmembrane transport
20/59 regulation of muscle system process
7/22 anion homeostasis
66/187 ion homeostasis
91/264 chemical homeostasis
59/166 cellular chemical homeostasis
4/10 platelet aggregation
16/30 inorganic anion transport
7/16 chloride transport
6/8 bicarbonate transport
7/14 sulfur compound transport
3/5 sulfate transport
11/33 detection of light stimulus
67/184 sensory perception
21/68 calcium ion transmembrane transport
29/95 divalent metal ion transport
14/24 larval locomotory behavior
58/142 locomotory behavior
15/34 larval behavior
12/33 male mating behavior
19/52 mating
9/25 courtship behavior
146/434 behavior
25/70 obsolete multi–organism behavior
18/46 rhythmic behavior
16/55 chemosensory behavior
6/11 L–glutamate transmembrane transport
13/24 L–amino acid transport
125/384 transmembrane transport
47/125 anion transmembrane transport
7/9 auditory behavior
32/73 organic acid transmembrane transport
50/125 organic anion transport
10/23 dicarboxylic acid transport
7/10 C4–dicarboxylate transport
8/23 fatty acid transmembrane transport
23/68 monocarboxylic acid transport
6/11 gamma–aminobutyric acid metabolic process
7/16 import across plasma membrane
20/43 import into cell
46/126 regulation of neurotransmitter levels
14/21 neurotransmitter uptake
100/344 multicellular organismal signaling
7/13 neutral amino acid transport
21/62 organic hydroxy compound transport
13/26 mechanosensory behavior
70/204 metal ion transport
27/47 sodium ion transport
8/28 potassium ion transport
85/284 cation transport
37/112 monovalent inorganic cation transport
14/24 vitamin transport
7/11 vitamin transmembrane transport
42/120 regulation of membrane potential
14/29 nerve development
17/26 positive regulation of nervous system process
60/170 organic hydroxy compound metabolic process
15/34 sterol metabolic process
28/74 steroid metabolic process
7/10 positive regulation of cholesterol transport
7/24 carbohydrate derivative transport
7/29 organophosphate ester transport
3/8 alditol phosphate metabolic process
5/12 heart jogging
16/52 embryonic heart tube morphogenesis
22/102 determination of bilateral symmetry
55/147 embryonic organ morphogenesis
15/43 ear morphogenesis
69/220 embryonic organ development
128/388 central nervous system development
8/14 heart valve development
39/126 stem cell differentiation
29/71 stem cell population maintenance
6/20 BMP signaling pathway
8/14 negative regulation of lipid metabolic process
5/17 negative regulation of protein binding
35/115 dendrite development
13/54 developmental cell growth
3/7 regulation of shoot system development
18/49 chaeta development
8/26 chaperone–mediated protein folding
9/17 heterochromatin organization
6/12 heterochromatin assembly
5/6 sulfur amino acid catabolic process
14/28 sulfur amino acid metabolic process
4/5 valine metabolic process
28/49 cellular amino acid catabolic process
7/7 erythrose 4–phosphate/phosphoenolpyruvate family amino acid catat
7/10 aromatic amino acid family catabolic process
37/82 carboxylic acid catabolic process

**Supplemental Figure 3**: Gene ontology (GO) enrichment of the "Biological Processes (BP)" category derived from *Acropora cervicornis* gene expression differences between the multistressor treatment (OAW+MP) relative to ambient conditions (AMB). Dendrograms depict sharing of genes between categories (categories with no branch length between them are subsets of each other), with the fractions corresponding to proportion of genes with an unadjusted $p < 0.05$ relative to the total number of genes within the category. Text size and boldness indicate the significance (Mann–Whitney *U* tests) of the term. Red categories are enriched in *A. cervicornis* in double stressor treatment (OAW+MP) while blue terms are underrepresented in OAW+MP treatment. The green square highlights the immunity-related GO terms that were used to identify immunity-related genes for heatmap visualization.

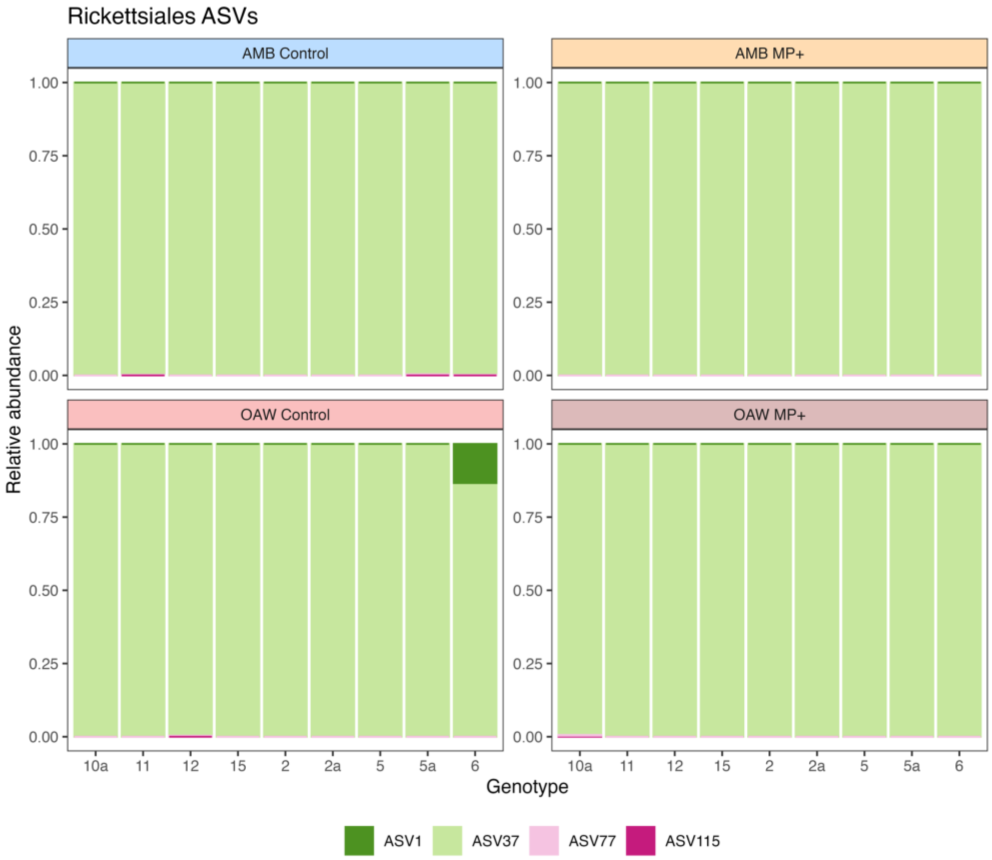


**Supplemental Figure 4**: Relative abundance of the four identified ASVs within order Rickettsiales across all samples. Samples were dominated by ASV 37, with background presence of the other 3 ASVs.

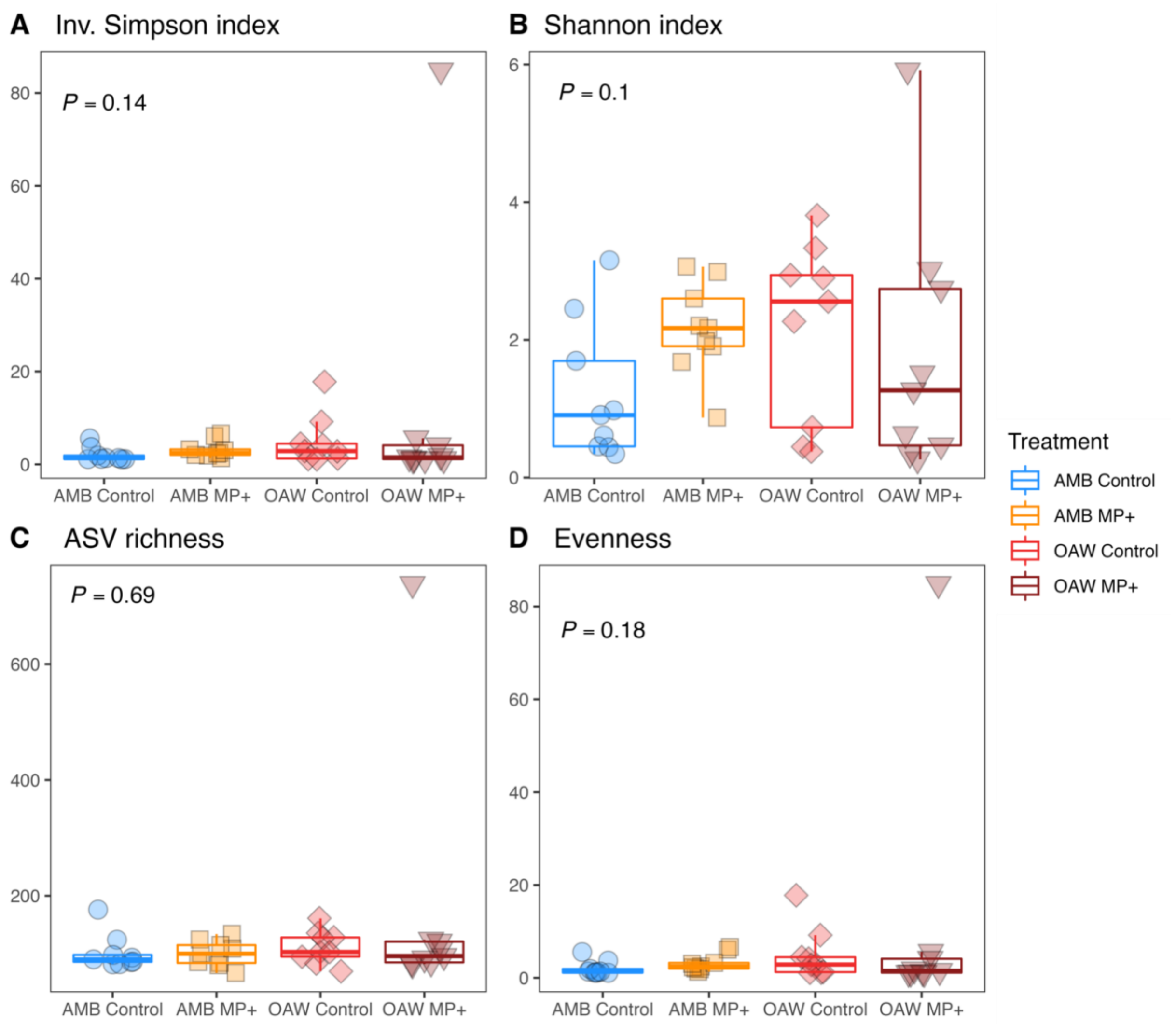


**Supplemental Figure 5**: Alpha diversity metrics across all samples in all treatments, depicting (A) Simpson index, (B) Shannon index, (C) species richness (ASVs), and (D) Evenness.

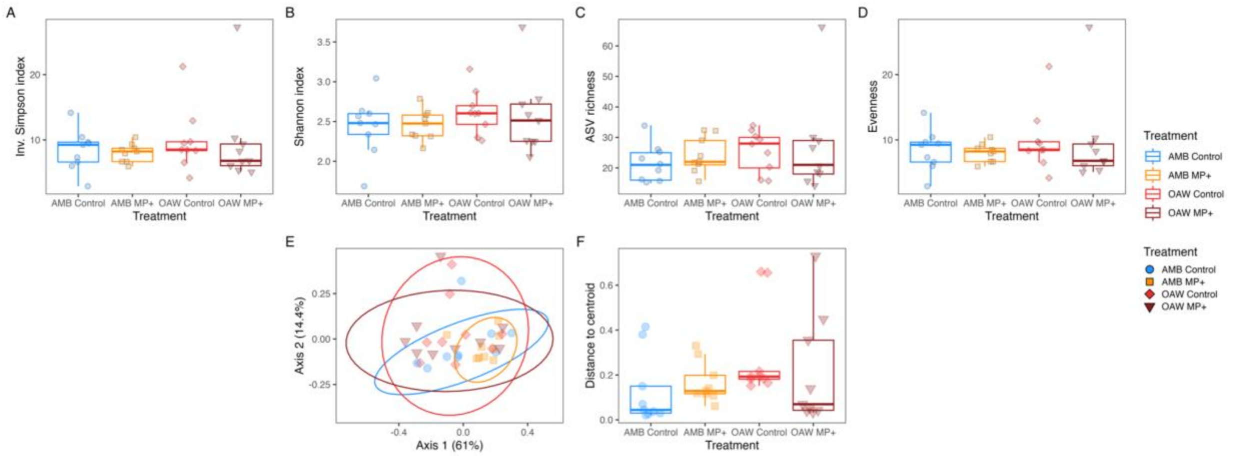


**Supplemental Figure 6**: Alpha (A-D) and beta (E-F) diversity metrics across all samples in all treatments once Aquarickettsia ASVs were removed. Plots depict (A) Simpson index, (B) Shannon index, (C) species richness, (D) Evenness, (E) multivariate ordination plots (PCoA) of between-sample Bray–Curtis dissimilarity of rarefied ASVs and (F) distance to treatment centroids. PCoA ellipses depict 95% confidence intervals and p-values indicate significance of treatment and genotype. Treatment is depicted by color in all panels: blue = ambient conditions (AMB), orange = ocean acidification and warming (OAW), red = microplastics (MP), and dark red = OAW and MP (OAW+MP).

### Supplemental Tables

**Supplemental Table 1**: Environmental conditions measured in every tank every day throughout the experiment by treatment and specific to each tank (temperature = °C; dissolved oxygen = ppm; salinity = ppt; pH = total scale).

| | Temperature | | | | Dissolved Oxygen | | | | Salinity | | | | pH | | | |
|---|---|---|---|---|---|---|---|---|---|---|---|---|---|---|---|---|
| | Mean | SD | Min | Max | Mean | SD | Min | Max | Mean | SD | Min | Max | Mean | SD | Min | Max |
| **Treatment** | | | | | | | | | | | | | | | | |
| AMB Control | 28.63 | 0.03 | 28.55 | 28.84 | 6.42 | 0.1 | 6.18 | 6.63 | 35.67 | 1.53 | 31.06 | 38.46 | 8.07 | 0.07 | 7.81 | 8.41 |
| AMB MP+ | 28.63 | 0.03 | 28.54 | 28.87 | 6.42 | 0.11 | 6.14 | 6.86 | 35.72 | 1.59 | 29.97 | 38.36 | 8.06 | 0.06 | 7.81 | 8.29 |
| OAW Control | 30.07 | 0.57 | 28.59 | 30.63 | 6.29 | 0.09 | 6.08 | 6.5 | 35.43 | 1.6 | 30.09 | 38.45 | 7.96 | 0.08 | 7.81 | 8.41 |
| OAW MP+ | 30.14 | 0.53 | 28.6 | 30.64 | 6.27 | 0.07 | 6.09 | 6.47 | 35.88 | 1.67 | 29.83 | 38.67 | 7.95 | 0.11 | 7.25 | 8.21 |
| **Tank** | | | | | | | | | | | | | | | | |
| 10aA - 1 | 30.2 | 0.54 | 28.63 | 30.63 | 6.23 | 0.06 | 6.11 | 6.32 | 37.01 | 0.75 | 35.29 | 37.91 | 7.96 | 0.08 | 7.8 | 8.12 |
| 10aB - 1 | 28.61 | 0.04 | 28.54 | 28.66 | 6.38 | 0.11 | 6.14 | 6.55 | 35.19 | 1.66 | 32.23 | 38.01 | 8.07 | 0.05 | 7.94 | 8.15 |
| 10aC - 1 | 30.14 | 0.55 | 28.6 | 30.6 | 6.31 | 0.1 | 6.16 | 6.48 | 34.62 | 1.4 | 31.49 | 37.35 | 7.95 | 0.06 | 7.85 | 8.08 |
| 10aD - 1 | 28.62 | 0.02 | 28.6 | 28.65 | 6.42 | 0.1 | 6.19 | 6.56 | 36.27 | 0.98 | 34.62 | 37.85 | 8.04 | 0.05 | 7.91 | 8.13 |
| 11A - 1 | 28.63 | 0.01 | 28.61 | 28.65 | 6.42 | 0.08 | 6.22 | 6.58 | 36.42 | 1.07 | 34.95 | 38.36 | 8.07 | 0.06 | 7.95 | 8.2 |
| 11B - 1 | 28.63 | 0.02 | 28.59 | 28.65 | 6.42 | 0.1 | 6.21 | 6.56 | 34.73 | 2.04 | 31.51 | 37.74 | 8.08 | 0.05 | 7.98 | 8.16 |
| 11E - 1 | 29.33 | 0.22 | 28.59 | 29.55 | 6.35 | 0.1 | 6.15 | 6.5 | 35.49 | 1.21 | 33.55 | 37.56 | 7.94 | 0.07 | 7.81 | 8.08 |
| 11E - 2 | 30.17 | 0.55 | 28.64 | 30.62 | 6.27 | 0.06 | 6.18 | 6.38 | 35.09 | 1.45 | 32.53 | 37.58 | 7.94 | 0.09 | 7.81 | 8.14 |
| 12A - 1 | 30.15 | 0.54 | 28.62 | 30.6 | 6.25 | 0.05 | 6.13 | 6.35 | 36.42 | 1.07 | 34.22 | 38.45 | 7.95 | 0.08 | 7.81 | 8.07 |
| 12B - 1 | 30.12 | 0.55 | 28.6 | 30.59 | 6.3 | 0.07 | 6.17 | 6.44 | 34.48 | 2.35 | 29.83 | 37.55 | 8.02 | 0.09 | 7.83 | 8.21 |
| 12E - 1 | 28.64 | 0.05 | 28.57 | 28.84 | 6.43 | 0.12 | 6.22 | 6.62 | 35.75 | 1.29 | 32.99 | 37.37 | 8.08 | 0.09 | 7.91 | 8.41 |
| 12E - 2 | 28.64 | 0.03 | 28.62 | 28.77 | 6.4 | 0.11 | 6.19 | 6.62 | 36.21 | 0.9 | 34.22 | 37.55 | 8.02 | 0.05 | 7.91 | 8.1 |
| 15C - 1 | 30.01 | 0.47 | 28.61 | 30.56 | 6.28 | 0.09 | 6.14 | 6.47 | 35.65 | 1.44 | 33.35 | 37.89 | 7.94 | 0.1 | 7.72 | 8.15 |
| 15E - 1 | 28.63 | 0.02 | 28.59 | 28.66 | 6.38 | 0.09 | 6.21 | 6.54 | 36.46 | 0.95 | 34.61 | 37.94 | 8.06 | 0.07 | 7.81 | 8.16 |
| 15E - 2 | 30.16 | 0.54 | 28.63 | 30.61 | 6.3 | 0.1 | 6.12 | 6.48 | 34.64 | 1.57 | 32.09 | 37.54 | 7.96 | 0.07 | 7.85 | 8.11 |
| 15E - 3 | 28.61 | 0.02 | 28.58 | 28.64 | 6.43 | 0.11 | 6.22 | 6.63 | 36.01 | 1.11 | 34.49 | 37.84 | 8.09 | 0.05 | 7.98 | 8.18 |
| 2A - 1 | 30.17 | 0.54 | 28.63 | 30.63 | 6.33 | 0.08 | 6.2 | 6.5 | 33.89 | 1.95 | 30.09 | 37.49 | 7.95 | 0.08 | 7.84 | 8.15 |
| 2aB - 1 | 30.16 | 0.54 | 28.6 | 30.62 | 6.25 | 0.06 | 6.15 | 6.34 | 36.68 | 0.99 | 34.84 | 38.25 | 7.87 | 0.22 | 7.25 | 8.05 |
| 2aC - 1 | 28.63 | 0.02 | 28.61 | 28.65 | 6.44 | 0.1 | 6.21 | 6.6 | 35.62 | 1.4 | 32.55 | 37.63 | 8.05 | 0.06 | 7.95 | 8.22 |
| 2aD - 1 | 28.63 | 0.02 | 28.55 | 28.66 | 6.42 | 0.09 | 6.24 | 6.56 | 36.1 | 1.4 | 33.51 | 38.46 | 8.08 | 0.07 | 7.82 | 8.17 |
| 2aE - 1 | 30.15 | 0.55 | 28.63 | 30.6 | 6.25 | 0.05 | 6.18 | 6.35 | 36.37 | 0.83 | 34.41 | 37.45 | 7.99 | 0.12 | 7.82 | 8.41 |
| 2B - 1 | 28.64 | 0.01 | 28.61 | 28.66 | 6.4 | 0.11 | 6.22 | 6.59 | 35.87 | 0.99 | 34.14 | 37.47 | 8.01 | 0.05 | 7.86 | 8.09 |
| 2D - 1 | 30.17 | 0.53 | 28.64 | 30.62 | 6.26 | 0.06 | 6.15 | 6.41 | 36.55 | 1.02 | 34.55 | 38.43 | 7.95 | 0.06 | 7.83 | 8.07 |
| 2E - 1 | 28.66 | 0.05 | 28.61 | 28.87 | 6.47 | 0.14 | 6.24 | 6.86 | 35.29 | 1.51 | 32.08 | 37.22 | 8.06 | 0.05 | 7.94 | 8.12 |
| 5aA - 1 | 30.15 | 0.56 | 28.63 | 30.62 | 6.3 | 0.06 | 6.2 | 6.42 | 35.54 | 1 | 33.55 | 36.98 | 7.96 | 0.07 | 7.83 | 8.09 |
| 5aC - 1 | 30.17 | 0.54 | 28.59 | 30.63 | 6.27 | 0.09 | 6.08 | 6.48 | 35.49 | 1.22 | 34.01 | 37.87 | 7.97 | 0.08 | 7.82 | 8.12 |
| 5aC - 2 | 28.64 | 0.02 | 28.6 | 28.67 | 6.42 | 0.11 | 6.18 | 6.63 | 35.7 | 1.47 | 32.62 | 38.18 | 8.03 | 0.07 | 7.81 | 8.19 |
| 5aE - 1 | 28.61 | 0.02 | 28.56 | 28.64 | 6.41 | 0.09 | 6.2 | 6.52 | 36.13 | 1.07 | 33.96 | 38.09 | 8.09 | 0.07 | 7.96 | 8.29 |
| 5B - 1 | 30.16 | 0.54 | 28.62 | 30.61 | 6.26 | 0.08 | 6.13 | 6.39 | 36.16 | 1.46 | 33.78 | 38.25 | 7.99 | 0.07 | 7.88 | 8.12 |
| 5C - 1 | 28.62 | 0.02 | 28.59 | 28.65 | 6.39 | 0.07 | 6.21 | 6.52 | 36.56 | 0.87 | 35.01 | 37.92 | 8.04 | 0.06 | 7.93 | 8.18 |
| 5C - 2 | 30.16 | 0.54 | 28.6 | 30.64 | 6.31 | 0.06 | 6.16 | 6.42 | 34.91 | 2.07 | 31.59 | 37.72 | 7.94 | 0.08 | 7.83 | 8.09 |
| 5D - 1 | 28.62 | 0.02 | 28.56 | 28.64 | 6.46 | 0.09 | 6.25 | 6.6 | 34.05 | 1.6 | 31.06 | 37.39 | 8.08 | 0.05 | 7.89 | 8.15 |
| 6B - 1 | 28.64 | 0.02 | 28.59 | 28.71 | 6.49 | 0.12 | 6.23 | 6.64 | 33.6 | 2.14 | 29.97 | 37.55 | 8.09 | 0.04 | 7.97 | 8.15 |
| 6B - 2 | 30.13 | 0.55 | 28.62 | 30.62 | 6.21 | 0.07 | 6.09 | 6.37 | 37.05 | 0.84 | 35.89 | 38.67 | 7.94 | 0.07 | 7.84 | 8.06 |
| 6D - 1 | 28.62 | 0.02 | 28.59 | 28.65 | 6.38 | 0.08 | 6.23 | 6.53 | 36.56 | 0.9 | 34.93 | 38.02 | 8.08 | 0.06 | 7.93 | 8.22 |
| 6E - 1 | 30.16 | 0.54 | 28.62 | 30.62 | 6.29 | 0.06 | 6.19 | 6.38 | 35.75 | 1.51 | 33.03 | 38.4 | 7.96 | 0.07 | 7.86 | 8.08 |

**Supplemental Table 2**: Metadata for all experimental *Acropora cervicornis* samples for gene expression analyses including sample ID, coral genotype (via SNPs), experimental treatment, raw reads, trimmed reads and reads that mapped to the *Acropora millepora* genome. Samples marked with ^ were removed from the dataset as they were clones within samples. Samples marked with * were removed due to low read count.

| Sample ID | Genotype | Treatment | Raw reads | trimmed | mapped |
|---|---|---|---|---|---|
| 2 | 10aA - 1 | OAW+MP | 5570866 | 2373894 | 688099 |
| 3 | 5aC - 2 | AMB | 5085548 | 2439160 | 818571 |
| 6 | 10aB - 1 | MP | 5426419 | 1956485 | 685167 |
| 8 | 6D - 1 | AMB | 6380098 | 2998772 | 966574 |
| 9 | 5aC - 1 | OAW | 4386666 | 1826327 | 674017 |
| 10 | 6B - 2 | OAW+MP | 5314015 | 2437133 | 834588 |
| 11 | 5B - 1 | OAW | 5816290 | 2453835 | 846094 |
| 12 | 5aE - 1 | MP | 4173771 | 1581695 | 609277 |
| ^14 | 12E - 1 | AMB | 5957546 | 2266810 | 828973 |
| 15 | 2E - 1 | MP | 4287069 | 2017225 | 648174 |
| 16 | 2B - 1 | AMB | 6580408 | 2819935 | 880982 |
| 17 | 6B - 1 | MP | 4731455 | 1354877 | 414581 |
| 19 | 10aC - 1 | OAW | 5426633 | 2536479 | 800716 |
| ^20 | 12E - 2 | MP | 5630212 | 2249677 | 863849 |
| 22 | 5aA - 1 | OAW+MP | 6403995 | 2632000 | 929270 |
| 23 | 5D - 1 | AMB | 4894064 | 2208318 | 780221 |
| ^24 | 12B - 1 | OAW+MP | 5530051 | 2322250 | 761130 |
| 25 | 5C - 1 | MP | 4858419 | 2005496 | 656062 |
| 26 | 10aD - 1 | AMB | 4419015 | 1643107 | 664303 |
| 28 | 5C - 2 | OAW+MP | 5079315 | 1585886 | 604654 |
| *29 | 2A - 1 | OAW | 5437415 | 625876 | 166828 |
| ^32 | 12A - 1 | OAW | 5469908 | 2594497 | 844935 |
| 34 | 2D - 1 | OAW+MP | 4970151 | 2423039 | 758436 |
| 35 | 6E - 1 | OAW | 5398937 | 1506896 | 521313 |

**Supplemental Table 3**: 16S reads through the *DADA2* filtering pipeline for all experimental fragments of *A. cervicornis*, including sample ID, genotype, and treatment condition.

| Sample | Genotype | Treatment | Input | Filtered | Denoised Forward | Denoised Reverse | Merged | Nonchimera |
|---|---|---|---|---|---|---|---|---|
| A10 | 6 | OAW MP+ | 25343 | 23991 | 23882 | 23969 | 23811 | 23811 |
| A11 | 5 | OAW Control | 27398 | 26006 | 25982 | 25971 | 25961 | 25961 |
| A12 | 5a | AMB MP+ | 45822 | 44152 | 44079 | 44127 | 43982 | 43982 |
| A2 | 10a | OAW MP+ | 99390 | 94836 | 94654 | 94676 | 94353 | 94205 |
| A3 | 5a | AMB Control | 77568 | 73174 | 73129 | 73130 | 73018 | 73018 |
| A6 | 10a | AMB MP+ | 44719 | 41897 | 41808 | 41848 | 41496 | 41496 |
| A8 | 6 | AMB Control | 50868 | 48504 | 48363 | 48414 | 48114 | 48073 |
| A9 | 5a | OAW Control | 33629 | 32048 | 31998 | 32015 | 31874 | 31842 |
| B10 | 5a | OAW MP+ | 26111 | 24910 | 24868 | 24838 | 24666 | 24641 |
| B11 | 5 | AMB Control | 36048 | 34782 | 34765 | 34754 | 34710 | 34710 |
| B3 | 2 | AMB MP+ | 36469 | 34374 | 34349 | 34353 | 34289 | 34289 |
| B4 | 2 | AMB Control | 93851 | 90460 | 90421 | 90323 | 90201 | 90201 |
| B5 | 6 | AMB MP+ | 24767 | 23566 | 23506 | 23540 | 23364 | 23328 |
| B7 | 10a | OAW Control | 114866 | 110660 | 110570 | 110599 | 110465 | 110452 |
| C1 | 5 | AMB MP+ | 29051 | 27927 | 27900 | 27896 | 27851 | 27851 |
| C10 | 2 | OAW MP+ | 75311 | 73356 | 73246 | 73255 | 72852 | 72852 |
| C11 | 6 | OAW Control | 24571 | 23594 | 23583 | 23578 | 23571 | 23571 |
| C2 | 10a | AMB Control | 26776 | 25590 | 25551 | 25561 | 25491 | 25491 |
| C4 | 5 | OAW MP+ | 43598 | 42054 | 42021 | 42021 | 41807 | 41723 |
| G5 | NA | blank_control | 1918 | 1795 | 1792 | 1792 | 1792 | 1792 |
| G6 | NA | blank_control | 169 | 162 | 153 | 158 | 153 | 153 |

**Supplemental Table 4**: Statistical model output of beta diversity assessments of all ASVs based on (A) treatment and (B) genotype.

A) ANOVA of all ASV dispersion

| | | Df | Sum Sq | Mean Sq | F value | Pr(>F) |
|---|---|---|---|---|---|---|
| **Treatment** | | | | | | |
| | Groups | 3 | 0.12 | 0.04 | 1.21 | 0.32 |
| | Residuals | 32 | 1.08 | 0.03 | | |
| **Genotype** | | | | | | |
| | Groups1 | 8 | 0.2 | 0.02 | 0.64 | 0.73 |
| | Residuals1 | 27 | 1.02 | 0.04 | | |

B) PERMANOVA of all ASVs

| | Df | Sum Of Sqs | $R^2$ | F value | Pr(>F) |
|---|---|---|---|---|---|
| Treatment | 3 | 0.23 | 0.09 | 1.05 | 0.42 |
| Genotype | 8 | 0.61 | 0.24 | 1.06 | 0.41 |
| *Residual* | *24* | *1.72* | *0.67* | | |
| *Total* | *35* | *2.55* | *1* | | |

**Supplemental Table 5**: Statistical model output of alpha diversity assessments of all ASVs.

| | Estimate | Std. Error | t value |
|---|---|---|---|
| **Simpson** | | | |
| (Intercept) | 0.67 | 0.1 | 6.79 |
| TreatmentAMB MP+ | -0.28 | 0.13 | -2.13 |
| TreatmentOAW Control | -0.23 | 0.13 | -1.79 |
| TreatmentOAW MP+ | -0.1 | 0.13 | -0.74 |
| **Shannon** | | | |
| (Intercept) | -0.1 | 0.28 | -0.35 |
| TreatmentAMB MP+ | 0.82 | 0.36 | 2.32 |
| TreatmentOAW Control | 0.58 | 0.36 | 1.63 |
| TreatmentOAW MP+ | 0.22 | 0.36 | 0.61 |
| **Richness** | | | |
| (Intercept) | 0.01 | 0 | 11.67 |
| TreatmentAMB MP+ | 0 | 0 | 0.08 |
| TreatmentOAW Control | 0 | 0 | -0.79 |
| TreatmentOAW MP+ | 0 | 0 | -1.01 |
| **Evenness** | | | |
| (Intercept) | 0.26 | 0.08 | 3.27 |
| TreatmentAMB MP+ | 0.21 | 0.11 | 2 |
| TreatmentOAW Control | 0.19 | 0.11 | 1.79 |
| TreatmentOAW MP+ | 0.09 | 0.11 | 0.83 |

**Supplemental Table 6**: Non-Aquarickettsia ASV beta (A-B) and alpha (C) diversity across samples.

**A) ANOVA of non Aquarickettsia dispersion**

| | Df | Sum Sq | Mean Sq | F value | P value |
|---|---|---|---|---|---|
| **Treatment** | | | | | |
| Groups | 3 | 0.09 | 0.03 | 2.02 | 0.13 |
| Residuals | 32 | 0.45 | 0.01 | | |
| **Genotype** | | | | | |
| Groups1 | 8 | 0.2 | 0.03 | 0.66 | 0.72 |
| Residuals1 | 27 | 1.04 | 0.04 | | |

**B) PERMANOVA of non Aquarickettsia**

| | Df | Sum Of Sqs | R2 | F | P value |
|---|---|---|---|---|---|
| Treatment | 3 | 0.55 | 0.11 | 1.29 | 0.11 |
| Genotype | 8 | 1.18 | 0.23 | 1.04 | 0.38 |
| *Residual* | *24* | *3.41* | *0.66* | | |
| *Total* | *35* | *5.14* | *1* | | |

**C) Alpha diversity metrics of non Aquarickettsia**

| | Estimate | Std. Error | t value | P value |
|---|---|---|---|---|
| **Simpson** | | | | |
| (Intercept) | 0.13 | 0.02 | 7.66 | 0 |
| TreatmentAMB MP+ | 0 | 0.02 | 0.11 | 0.91 |
| TreatmentOAW Control | -0.02 | 0.02 | -0.89 | 0.37 |
| TreatmentOAW MP+ | -0.02 | 0.02 | -1.12 | 0.26 |
| **Shannon** | | | | |
| (Intercept) | 1.55 | 0.03 | 46.19 | 0 |
| TreatmentAMB MP+ | 0.02 | 0.04 | 0.46 | 0.65 |
| TreatmentOAW Control | 0.05 | 0.04 | 1.24 | 0.23 |
| TreatmentOAW MP+ | 0.07 | 0.04 | 1.5 | 0.15 |
| **Richness** | | | | |
| (Intercept) | 3.02 | 0.1 | 28.93 | 0 |
| TreatmentAMB MP+ | 0.14 | 0.12 | 1.11 | 0.28 |
| TreatmentOAW Control | 0.19 | 0.12 | 1.51 | 0.15 |
| TreatmentOAW MP+ | 0.15 | 0.12 | 1.22 | 0.23 |
| **Evenness** | | | | |
| (Intercept) | 1.25 | 0.04 | 34.5 | 0 |
| TreatmentAMB MP+ | 0.03 | 0.05 | 0.57 | 0.57 |
| TreatmentOAW Control | -0.01 | 0.04 | -0.32 | 0.75 |
| TreatmentOAW MP+ | -0.04 | 0.04 | -0.94 | 0.34 |
